# Cooperative development of logical modelling standards and tools with CoLoMoTo

**DOI:** 10.1101/010504

**Authors:** Aurélien Naldi, Pedro T. Monteiro, Christoph Müssel, the Consortium for Logical Models and Tools, Hans A. Kestler, Denis Thieffry, Ioannis Xenarios, Julio Saez-Rodriguez, Tomas Helikar, Claudine Chaouiya

**Author notes:** The complete list of members of the Consortium for Logical Models and Tools is provided in the Acknowledgement section.

## Abstract

The identification of large regulatory and signalling networks involved in the control of crucial cellular processes calls for proper modelling approaches. Indeed, models can help elucidate properties of these networks, understand their behaviour, and provide (testable) predictions by performing in silico experiments. In this context, qualitative, logical frameworks have emerged as relevant approaches as demonstrated by a growing number of published models, along with new methodologies and software tools. This productive activity now requires a concerted effort to ensure model reusability and interoperability between tools. Here, we outline the logical modelling framework and present the most important achievements of the **Co**nsortium for **Lo**gical **Mo**dels and **To**ols, along with future objectives. This open community welcomes contributions from all researchers interested in logical modelling or in related mathematical and computational developments.

## 2 Motivation

The rapid development of novel biomolecular technologies has fostered the study of complex regulatory systems during the last decade. Mathematical models have become invaluable tools for understanding the dynamical behaviour of such systems. There are various types of formalisms, which differ in the level of detail and model complexity (de Jong, 2002; Karlebach and Shamir, 2008). Logical (or logic), discrete models comprise the most abstract dynamic models and constitute nowadays a popular modelling framework. In logical models, components are represented by discrete variables with a small range of possible values, with the most extreme case being Boolean models, where each component can be either active or inactive (Kauffman, 1969; Thomas and d’Ari, 1990). Regulatory effects are defined by logical rules or lookup tables. The relative simplicity of these models provides several advantages over more complex modelling formalisms, such as systems of differential equations. In particular, logical models do not require precise knowledge of kinetic parameters, which makes them suitable for large models comprising up to several hundreds of components. Due to their qualitative nature, it is possible to incorporate various kinds of information in logical models. For example, natural-language statements from publications or expert knowledge on regulatory interactions can easily be transformed into logical rules.

Although logical models provide a rough approximation of concentration levels, they can reproduce the behaviour of many biological systems as illustrated in examples of case studies below (for recent reviews see Bornholdt, 2005; Saadatpour and Albert, 2013). The discrete step functions resulting from simulations of logical models constitute a plausible simplification of typical sigmoidal response curves (de Jong, 2002). Biomolecular measurements can be analysed in the discrete framework by applying specialised discretisation procedures (Shmulevich and Zhang, 2002; Hopfensitz *et al.*, 2012).

Logical models have been successfully applied to a wide range of regulatory and signalling systems, in a variety of organisms. Examples include: yeast cell cycle (Li *et al.*, 2004; Orlando *et al.*, 2008; Davidich and Bornholdt, 2008; Fauré and Thieffry, 2009; Todd and Helikar, 2012), pathogen-host interactions (Madrahimov *et al.*, 2012; Franke *et al.*, 2008; Thakar *et al.*, 2007), development of the sea urchin embryo (Peter *et al.*, 2012), development of Drosophila melanogaster (Sánchez and Thieffry, 2001, 2003; Albert and Othmer, 2003; Sánchez *et al.*, 2008), mammalian cell cycle (Fauré *et al.*, 2006), murine cardiac development (Herrmann *et al.*, 2012), determination of cell fates in human (Calzone *et al.*, 2010; Grieco *et al.*, 2013), multi-scale signal transduction in human epithelial cells (Helikar *et al.*, 2013), human T-cell receptor signaling (Saez-Rodriguez *et al.*, 2007; Zhang *et al.*, 2008) and T-cell differentiation (Naldi *et al.*, 2010).

The *Consortium for Logical Models and Tools* (CoLoMoTo, http://colomoto.org) was informally launched during a meeting at Instituto Gulbenkian de Ciência (Portugal) in November 2010 to gather interested scientists and promote the cooperative development of shared standards and tools. This meeting was followed by a second one at the European Bioinformatics Institute (United Kingdom) in March 2012, which focused more specifically on a common standard for the exchange of logical model (SBML qual, see below). The third meeting was held at the University of Lausanne (Switzerland) in April 2014, bringing together 22 participants from 14 different institutions. It included sessions devoted to scientific presentations covering methodological and computational developments, as well as models for real case applications. In addition, the participants discussed several topics requiring community consensus, including the development of standards, collaborative tools, and model repositories.

CoLoMoTo is an international open community that brings together modellers, curators, and developers of methods and tools. It aims at the definition of standards for model representation and interchange, and the establishment of criteria for the comparison of methods, models and tools. Finally, CoLoMoTo seeks to promote these methods, tools and models.

Here, we briefly introduce the logical modelling approach. We also summarise the recent achievements of the consortium, as well as the outcomes of the last meeting regarding future directions.

## 3 Logical modelling

This section presents a brief overview of the logical framework diversity, providing main references for further information.

### 3.1 Logical models

The development of a logical model for a regulatory and/or signalling network usually starts with the definition of a graph encompassing relevant regulatory components (nodes or vertices) and their interactions (edges or arcs). A discrete variable is associated with each component to denote its concentration or activity level. A logical function (or logical rule) defines the evolution (target value of the corresponding variable) of each component, depending on the levels of its regulators. One can distinguish Boolean models, where all variables are Boolean (0 or 1), (Kauffman, 1969; Thomas, 1973; Bornholdt, 2005), from multi-valued logical models, where variables can take additional values (Thomas, 1991). Regarding the logical rules, some models are limited to specific classes of functions, such as threshold functions (Li *et al.*, 2004; Bornholdt, 2008) or canalysing functions (Kauffman *et al.*, 2003). Logical functions can be randomly selected (Shmulevich *et al.*, 2002) according to predefined specifications. In *Random Boolean Networks*, all interactions and regulatory functions are randomly assigned. They can be utilised to study the global behaviour of certain classes of networks.

For multi-cellular systems as in developmental processes (e.g. epithelial pattern formation), composition methods have been proposed to define models in terms of intracellular networks inter-connected through integration functions accounting for cell-cell communication (Mendes *et al.*, 2013; Fauré *et al.*, 2014).

### 3.2 Logical states and simulations

The current state of a logical model is encoded in a vector giving the levels of all components. As the number of integer values allowed for each variable is finite (and usually small), the state space is also finite. The evolution of each component at a given state is determined by its logical function but may also depends on the updating policy. With the *synchronous* policy, all components are updated simultaneously, leading to a deterministic behaviour, where each state has at most one successor. With the *asynchronous* policy, only one component is updated at a given time, i.e. a state has as many successors as the number of components called to update. More sophisticated updating policies have been proposed, which can be deterministic (e.g. synchronous, block-sequential Robert (1986); Aracena *et al.* (2013)), yield multiple successors (e.g. asynchronous, priority classes (as in Fauré *et al.*, 2006), or introduce stochastic choices (e.g. through the random selection of a single successor from those enabled by the asynchronous policy (Harvey and Bossomaier, 1997). An exhaustive list of these policies is beyond our scope, but it is important to realise that model behaviour may substantially change with the choice of updating.

The dynamics of a logical model can be conveniently represented by a State Transition Graph (STG), where nodes denote states and edges denote enabled state transitions. Each transition may involve the update of one or more components, as determined by the logical regulatory functions and the updating policy.

#### Logical framework extensions

Briefly described below are a few of the extensions that have been proposed to extend the scope of the formalism.

Time delays were an early concern of R. Thomas (see Thomas and d’Ari (1990), and Thomas (2013) for a recent discussion). As a matter of fact, several extensions of the logical framework intend to integrate kinetic information. For example, Siebert and Bockmayr (2006) proposed to consider constraints on time delays associated with variables updates. Boolean models can also be extended by associating discrete time delays with the inputs of the transition functions, hence incorporating asynchronism in a synchronous and deterministic updating policy (Helikar and Rogers, 2009; Müssel *et al.*, 2014). Furthermore, predicates that depend on absolute time points can be incorporated to model changes in the modelled system over time, e.g. the influence of external factors or processes like cell differentiation. In Peter *et al.* (2012), temporal and spatial constraints are simply expressed as additional logical rules.

In classical Boolean network simulations, external components (input nodes with no regulator) are fixed at 0 or 1 for the duration of the simulation. Alternatively, the values of these external components can be set according to a probabilistic distribution over time to cope with variable environmental conditions. Furthermore, the activity levels of individual model components can be defined as the ratio of 0’s and 1’s over a number of time steps and thereby enable analyses mimicking experimental dose-response curves (Helikar *et al.*, 2012b).

Finally, logical models can be refined and translated into continuous models with similar dynamical properties by generating standardised Ordinary Differential Equations (ODEs) based on the logical rules (Di Cara *et al.*, 2007; Krumsiek *et al.*, 2010; Ouattara *et al.*, 2010). Such continuous models introduce numerous parameters, which then need to be defined or estimated. Alternatively, Piecewise-Linear Differential Equations (PLDEs) (Glass and Kauffman, 1973) approximate the switch-like properties of regulatory interactions. PLDE can be simulated using the Genetic Network Analyzer (de Jong *et al.*, 2003), provided that qualitative relationships between parameters are defined.

### 3.3 Analysis of logical models

A first important property of a logical model is the repertoire of its asymptotic behaviours, namely its set of *attractors*. Attractors are sets of states from which the system cannot escape. They are often assumed to denote biologically relevant behaviours, e.g. Hopfensitz *et al.* (2013). For example, *stable states* may describe cell fates such as differentiation or apoptosis, whereas *complex attractors* may represent oscillatory properties, e.g. cell division cycle or circadian rhythms (Huang, 1999). Alternative stable states have been associated with multiple differentiated T-helper subtypes (Naldi *et al.*, 2010), with specific expression patterns associated with the different segments of fruit fly embryo (Albert and Othmer, 2003), with different cardiac progenitor cells (Herrmann *et al.*, 2012), and with cell fates in response to death receptor engagement (Calzone *et al.*, 2010). In contrast, Fauré *et al.* (2006) associated a complex cyclic attractor with the oscillatory behaviour of the mammalian cell cycle.

Reachability properties are also of high interest. They represent the capability of the model to generate a particular trajectory (a path in the STG). For example, one can analyze the model to identify a trajectory from an initial condition toward a specific attractor.

#### Dynamical analysis by simulation

The study of the dynamical properties can be performed through the analysis of the STG reachable from a set of initial states (or all possible states for the complete STG). In the STG, an attractor corresponds to a terminal Strongly Connected Component (SCC), defined as a set of states such that each state can be reached from all other states with no transition leaving this set of states. Whenever a terminal SCC encompasses a unique state, the attractor is a stable state, whereas a SCC containing at least two states denotes a cyclic attractor. To ease the identification of attractors and the analysis of the STG structure, more compact representations can be generated, including strongly connected component graphs, or more compressed hierarchical transition graphs (Bérenguier *et al.*, 2013).

STG analysis rapidly becomes intractable when the size of the model increases. The use of efficient data structures to represent the dynamics greatly facilitates the identification of the attractors in Boolean models (Garg *et al.*, 2008).

To avoid the combinatorial explosion of the state space, Stoll *et al.* (2012) proposed a Boolean Kinetic Monte-Carlo algorithm which applies a continuous time Markov process to a Boolean state space. By assigning a transition rate to each component and considering time as a real number, it uses Monte-Carlo simulations to compute the temporal evolution of probability distributions and to estimate stationary distribution of logical states.

The verification of reachability properties quickly becomes time consuming even for small models, calling for semi-automated methods. One popular approach is *model checking* (Clarke *et al.*, 2000), which has been widely used for the verification of software and hardware systems during the last 30 years. More recently, model checking has been successfully applied to logical models (Bernot *et al.*, 2004; Monteiro and Chaouiya, 2012), as well to piecewise-linear differential models (Batt *et al.*, 2005), Petri nets (Gilbert *et al.*, 2007), among others.

#### Static analysis from model structure

Some *static analysis* methods were proposed to mitigate combinatorial explosion, i.e. by deducing dynamical information from the model itself, rather than computing explicit state transitions. In this context, the first question of interest is the identification of attractors, which is currently the subject of intense research.

The simplest attractors consists in states for which no component can be updated, called *stable states*. They are independent of the updating policy, have a simple formal definition and can be efficiently computed using constraint-solving methods (Devloo *et al.*, 2003; Naldi *et al.*, 2007) or polynomial algebra (Veliz-Cuba *et al.*, 2010). The notion of stability has been generalised from single states to sub-spaces of states Siebert (2011) with applications in model reduction and attractor detection. Like stable states, stable subspaces are independent of the updating policy and can be computed by constraint-solving methods Klarner *et al.* (2014).

In contrast, complex attractors depend on the updating policy and are more tricky to compute, in particular in the asynchronous case. A number of methods have been proposed recently for their identification, including an efficient SAT-based algorithm for the synchronous case (Dubrova and Teslenko, 2011). Turning to the asynchronous case, a method has been recently proposed to circumscribe regions of the state space that contain complex attractors by identifying the nodes whose state stabilises in these attractors (Zañudo and Albert, 2013).

Multi-stability (existence of multiple attractors) or sustained oscillations (or homeostasis) require the presence of positive or negative regulatory circuits in the corresponding regulatory graph (Thomas, 1981; Thieffry, 2007; Remy *et al.*, 2008). The connection between regulatory graph structure and dynamics is currently studied by several groups, and can be used to deduce dynamical properties, at least in some specific cases (Comet *et al.*, 2013). Attractors reachability from specific initial conditions can also be studied through static analysis in specific cases (Paulevé *et al.*, 2012).

Finally, model reduction techniques aim at building a simpler model preserving most dynamical properties. In this respect, automated reduction methods facilitate this process by properly rewriting logical rules (Naldi *et al.*, 2011).

## 4 Standard and tools

While numerous independent software packages supporting logical models have been proposed over the years, no standard existed for exchanging models between them. The CoLoMoTo consortium was created to foster the design of such standards. Collaborative efforts within CoLoMoTo already led to a novel model exchange format, SBML qual, along with a standard Java^TM^ LogicalModel library to handle logical models.

### SBML qual

(Qualitative Models package for SBML) is an extension of the Systems Biology Markup Language (SBML) Level 3 standard (Hucka *et al.*, 2010). It is designed for the representation of multi-valued qualitative models of biological networks (Chaouiya *et al.*, 2013a,b). After various meetings and refinements by logical modelling software developers, this new package was accepted by the SBML community in 2011 and finally approved by the SBML Editors in the spring of 2013. The SBML qual extension is supported in libSBML (Bornstein *et al.*, 2008) and JSBML (Dräger *et al.*, 2011). Models encoded in SBML qual can be submitted to the BioModels database, which also includes a branch dedicated to non-metabolic models automatically generated from pathway resources (Chelliah *et al.*, 2013; Büchel *et al.*, 2013).

The SBML qual format supports the definition of the model itself as a list of components (or SBML *species*), each with a maximum value and an optional initial value. Interactions between these components and logical rules complete the model definition. This first version thus accounts for generic multi-valued logical models. The extensions briefly described in Section 3 and simulation settings (e.g. updating mode, perturbations, etc.) are currently not yet supported. It is the goal of the consortium to tackle these issues.

### The LogicalModel library

provides a reference implementation of logical models as supported by SBML qual, relying on logical functions and decision diagrams. While SBML qual is the main supported format, the API enables the definition of additional import and export formats. The LogicalModel library can thus act as a format conversion module for tools that do not support SBML qual directly. It is freely available at https://github.com/colomoto/logicalmodel.

We now shortly describe tools that support the new SBML format, either directly or through the LogicalModel library.

### BoolNet

is a R package for the construction, analysis and simulation of different types of Boolean networks (Müssel *et al.*, 2010), freely available from the R project at http://cran.r-project.org/. It supports synchronous, asynchronous and probabilistic updatings, as well as time delays and temporal predicates as discussed in Section 3. State transition graphs, sequences and attractors can be analyzed and visualised. BoolNet further provides tools to assess model robustness, as well as to compare a given model with populations of randomly generated models. The package also comprises reverse-engineering methods enabling the inference of models from measurements. Finally, it can import and export models in the SBML qual format.

### Cell Collective

(http://www.thecellcollective.org) is a web-based platform for the construction, simulation, and analysis of Boolean-based models (Helikar *et al.*, 2012b). Simulations in Cell Collective can be done in a real-time, interactive fashion. Using extensions discussed in Section 3.2, simulation output results can be also represented as dose-response curves. The platform’s Bio-Logic Builder tool (Helikar *et al.*, 2012a) enables models to be constructed and edited in terms of biochemical/biological regulatory mechanism, avoiding explicit (error-prone) manual introduction of the Boolean functions. Logical modelling thereby becomes more accessible to wet lab biologists. The platform includes a knowledge base facilitating collaborative model annotation, in order to track the evidence associated with each interaction. The objective of this knowledge base is to enable users to create intuitable models and make them more appealing for the community to review, use, and expand. Cell Collective further provides a repository of Boolean models, which can be directly simulated and analysed (currently over 30 published models are available). Shared on the web, these models can also be exported as SBML qual files (or as text files listing logical expressions and truth tables).

### CellNetAnalyzer

(http://www.mpi-magdeburg.mpg.de/projects/cna/) (CNA) is a MATLAB toolbox providing a graphical user interface with various computational methods and algorithms for exploring structural and functional properties of metabolic, signalling, and regulatory networks (Klamt *et al.*, 2007). Whereas metabolic networks are formalised and analysed by stoichiometric modelling techniques, signal transduction and (gene) regulatory networks are represented in CNA as logical networks (both Boolean and multi-valued logic are supported). Among other features, CNA supports logical steady state analysis (e.g., for studying the input-output behaviour of signalling networks), the computation of minimal intervention sets (to enforce or block certain behaviors) (Samaga *et al.*, 2010), and, via the ODEfy plugin (Krumsiek *et al.*, 2010), discrete simulations of Boolean models, as well as simulation of logic-based ODEs derived from Boolean models. Furthermore, methods for studying properties of interaction (or regulatory) graphs underlying logical models are provided such as enumeration and analysis of feedback loops and signalling paths, and analysis of global interdependencies (Klamt *et al.*, 2006). The algorithms can be used within the GUI or via API functions, and SBML qual export is supported.

### CellNOpt

(http://www.cellnopt.org) is a free open-source software dedicated to train logical models to experimental data (Terfve *et al.*, 2012). The core of CellNOpt consists of a set of R packages, but can also be used through a Python wrapper, as well as a Cytoscape plug-in (CytoCopteR), which provides support for the SBML qual format, and a bridge for other Cytoscape plugins (Saito *et al.*, 2012). Starting with a Prior Knowledge Network (i.e. a network topology), CellNOpt generates a family of optimal models by minimising the difference between data and simulation results and penalising model size. Model optimisation can be performed using a variety of formalisms: (i) Boolean models, using synchronous updating or steady-state computation, (ii) semi-quantitative constrained Fuzzy logic, (iii) ordinary differential equations (ODEs) derived from the logical model. CellNOpt is integrated with data-driven reverse engineering methods via the CNORFeeder package (Eduati *et al.*, 2012).

### GINsim

(http://ginsim.org) is a Java application for the construction and analysis of multi-valued logical models (Naldi *et al.*, 2009). GINsim provides a user-friendly graphical interface to define the model, from the delineation of the regulatory graph to the definition of logical rules and simulation parameters. GINsim supports the simulation of logical models by generating state transition graphs for synchronous and asynchronous updatings, as well as priority classes. In addition, GINsim offers efficient methods for the reduction and composition of models, the determination of stable states (without exploring the STG), as well as for the analysis of regulatory circuits associated with crucial dynamical properties. Finally, GINsim includes a collection of import and export filters to enable the exchange of models with other tools, including SBML qual, as well as several Petri net and model-checker formats.

### MaBoSS

(http://maboss.curie.fr/) is a C++ software for the simulation of continuous time Markov processes directly derived from Boolean models (Stoll *et al.*, 2012). Using a specific language to define the model and transition rates, it applies the Gillespie algorithm to compute temporal trajectories. It estimates time dependent probabilities of logical model state. In addition, asymptotic behaviours (e.g. probability of an attractor) are computed. The LogicalModel library provides an export filter to the MaBoSS format, while a conversion of MaBoSS models into SBML qual is planned.

### EpiLog

(http://ginsim.org/epilog) is a Java software for the qualitative simulation of epithelial patterning. It defines an epithelium as a 2D grid of hexagonal cells, each containing a logical model of its regulatory network. External influences from neighbouring cells and from the environment are integrated as input components of each cell. The software provides a graphical user interface to ease the definition of initial conditions and perturbations, and to visualise simulations directly on the grid of cells. It uses the LogicalModel library to load the logical model(s) (in SBML qual format) and to evaluate the regulatory functions. Simulations combine a/synchronous updating components within each cell, with synchronous updating of identical components across cells.

### SQUAD and BoolSim

(http://www.vital-it.ch/software/SQUAD/ and (http://www.vital-it.ch/software/genYsis, a wrapper around BoolSim). BoolSim implements an efficient algorithm, based on binary decision diagrams, to identify all attractors (stable states as well as cyclic attractors) of Boolean models. SQUAD derives continuous dynamical models from logical models, and performs continuous simulations, allowing the modification of several parameters (Di Cara *et al.*, 2007). Starting with the discrete steady states identified by BoolSim, it localises the corresponding continuous steady states. The LogicalModel library provides import and export filters for the BoolSim format.

## 5 Conclusions and prospects

Logical modelling of biological regulatory networks constitutes a very active field involving scientists with diverse interests, ranging from methodological developments and computational implementations to real case applications. With the aim to foster synergies between these multiples developments, the Consortium for the development of Logical Models and Tools (CoLoMoTo) was launched four years ago and already delivered the following results: (1) the definition of the SBML Level 3 Qualitative Model (SBML qual) package for the representation of multi-valued qualitative models of biological networks; (2) the implementation of the standard LogicalModel library, which can be used by various modelling and simulation tools.

During the last meeting in April 2014, is was decided to organise CoLoMoTo activities along four main axes.

The first axis aims at standardisation. The reproducibility of results is enforced by defining and extending standards for the representation and interchange of models and their simulation parameters. Useful enhancements of SBML qual have already been discussed by the community. Improvements considered include the definition of models where rules are not (all) instantiated, models for which timing constraints (or rates) are specified, etc. In addition, further integration with SBML Core concepts will be needed. In particular, such integration would facilitate support of hybrid models, which combine features of both discrete and continuous formalisms. This activity is developed in close connection with the COMBINE (COmputational Modeling in BIology NEtwork) community, whose raison d’être is the development of standards for model interchange. One future direction addresses the issue of exchanging simulation settings for logical models, using the Simulation Experiment Description Markup Language (SED-ML) (Waltemath *et al.*, 2011). Moreover, the adoption of the Kinetic Simulation Algorithm Ontology (KiSAO) will permit a better description of the algorithms, their parameters and relationships (Courtot *et al.*, 2011). Finally, CoLoMoTo is currently working on the definition of a controlled vocabulary (ontology) covering the essential terms related to logical modelling, with textual definitions and corresponding references.

The second axis aims at defining an umbrella model repository with links to existing model repositories. Authors of manuscripts describing new logical models will be encouraged to publish their models in one of these repositories.

The third axis consists in defining benchmarks for the comparison of models and tools. Furthermore, successful modelling collaborations between experimentalists and modellers will be documented, in order to guide novel projects.

The fourth axis is the creation of a repository of methods and tools that are made available by the different research groups working on logical modelling. This repository should not only list the different features and functionalities provided by each of the tools and methods, but also provide guidelines for the selection of tools and methods suitable for typical use-cases.

Recently launched (http://colomoto.org), a multi-purpose umbrella portal provides access to the reports of the CoLoMoTo meetings, along with contact data for all involved research groups. An important current aim consists in reaching more scientists and making them aware of existing models, methods and tools, which could be used for their own research lines. We thus conclude this letter by warmly inviting the international community interested in logical modelling to participate in future CoLoMoTo activities.

## Acknowledgement

Members of the Consortium for Logical Models and Tools: Reka Albert, Matteo Barberis, Laurence Calzone, Claudine Chaouiya, Anastasia Chasapi, Thomas Cokelaer, Isaac Crespo, Julien Dorier, Andreas Dräger, Tomas Helikar, Céline Hernandez, Hidde de Jong, Sarah M. Keating, Hans A. Kestler, Steffen Klamt, Hannes Klarner, Reinhard Laubenbacher, Nicolas Le Novère, Pedro T. Monteiro, Christoph Müssel, Aurélien Naldi, Anne Niknejad, Nicolas Rodriguez, Julio Saez-Rodriguez, Heike Siebert, Denis Thieffry, Ioannis Xenarios, Jorge G. T. Zañuudo.

### Funding

We acknowledge the financial support from the Fundaç ão Calouste Gulbenkian, the European Commission FP6 NoEs ENFIN LSHG-CT-2005-518254, the European Bioinformatics Institute (EMBL-EBI), the European Union through the BioPreDyn project to JSR (ECFP7-KBBE-2011-5 grant No 289434), Swiss State Secretariat for Research and Innovation, AgeBrainSysBio contract No 305299 and SystemsX.ch - Swiss Systems Biology Initiative which made the three CoLoMoTo meetings possible.

